# A post-translational modification of the sheath modulates *Francisella* type VI secretion system assembly and function

**DOI:** 10.1101/370957

**Authors:** Jason Ziveri, Cerina Chhuon, Anne Jamet, Guénolé Prigent, Héloïse Rytter, Fabiola Tros, Monique Barel, Mathieu Coureuil, Claire Lays, Thomas Henry, Nicholas H Keep, Ida Chiara Guerrera, Alain Charbit

**Affiliations:** Université Paris Descartes, Sorbonne Paris Cité, Bâtiment Leriche, Paris, INSERM U1151 - CNRS UMR 8253, Institut Necker-Enfants Malades. Team 11: Pathogenesis of Systemic Infections, Paris, France; Plateforme protéomique 3P5-Necker, Université Paris Descartes - Structure Fédérative de Recherche Necker, INSERM US24/CNRS UMS3633, Paris 75014, France; Crystallography, Institute for Structural and Molecular Biology, Department of Biological Sciences Birkbeck, University of London, United Kingdom; CIRI, Centre International de Recherche en Infectiologie, Université Lyon, Inserm, U1111, University Claude Bernard Lyon 1, CNRS, UMR5308, École Normale Supérieure de Lyon, Labex Ecofect, Eco-evolutionary dynamics of infectious diseases, F-69007, LYON, France

**Author notes:** Corresponding authors: Ida Chiara Guerrera, Alain Charbit.; —. Bâtiment Leriche. 14 Rue Maria Helena Vieira Da Silva CS 61431 - 75993 PARIS – FRANCE.

**Keywords:** Type 6 secretion system, Phosphoproteome, *Francisella tularensis*, IglB.

## Abstract

*Francisella tularensis* is a facultative intracellular pathogen that causes the zoonotic disease tularemia in human and animal hosts. This bacterium possesses a non-canonical type VI secretion systems (T6SS) required for phagosomal escape and access to its replicative niche in the cytosol of infected macrophages. KCl stimulation has been previously used to trigger assembly and secretion of the Francisella T6SS in culture. We found that the amounts of essentially all the TSS6 proteins remained unchanged upon KCl stimulation. We therefore hypothesized that a post-translational modification might be involved in T6SS assembly. A whole cell phosphoproteomic analysis allowed us to identify a unique phosphorylation site on IglB, the TssC homologue and key component of the T6SS sheath. Importantly, the phosphorylated form of IglB was not present in the contracted sheath and 3D modeling indicated that the charge repulsion provoked by addition of a phosphogroup on tyrosine 139 was likely to weaken the stability of the sheath structure. Substitutions of the phosphorylatable residue of IglB (tyrosine 139) with alanine or with phosphomimetics prevented T6SS formation and totally impaired phagosomal escape. In contrast, the substitution with the non-phosphorylatable aromatic analog phenylalanine impaired but did not prevent phagosomal escape and cytosolic bacterial multiplication in J774-1 macrophages. Altogether these data suggest that phosphorylation of the sheath participates to T6SS disassembly. Post-translational modifications of the sheath may represent a previously unrecognized mechanism to finely modulate the dynamics of T6SS assembly-disassembly.

Data are available via ProteomeXchange with identifier PXD012507.

**Synopsis:** *Francisella* possesses a non-canonical T6SS that is essential for efficient phagosomal escape and access to the cytosol of infected macrophages. KCl stimulation has been previously used to trigger assembly and secretion of the Francisella T6SS in culture. We found that KCl stimulation did not result in an increased production of TSS6 proteins. We therefore hypothesized that a post-translational modification might be involved in T6SS assembly. Using a global and site-specific phosphoproteomic analysis of *Francisella* we identified a unique phosphorylation site on IglB, the TssC homologue and a key component of the T6SS contractile sheath. We show that this site plays a critical role in T6SS biogenesis and propose that phosphorylation may represent a new mechanism affecting the dynamics of sheath formation.

## Introduction

*Francisella tularensis* is the causative agent of the zoonotic disease tularemia (Luque-Larena, Mougeot et al., 2017, Maurin & Gyuranecz, 2016, Sjostedt, 2007, Sjostedt, 2011). This facultative intracellular pathogen is able to infect a variety of different cell types but, *in vivo*, is thought to replicate and disseminate mainly in phagocytes (Santic, Molmeret et al., 2006). Four major subspecies (subsp) of *F. tularensis* are currently listed that all cause a fulminant disease in mice that is similar to tularemia in human (Kingry & Petersen, 2014, McLendon, Apicella et al., 2006). Although the subsp *novicida* (also called *F. novicida) is* rarely pathogenic in human, its genome shares a high degree of nucleotide sequence conservation with the human pathogenic species and is thus widely used as a model organism (Brodmann, Dreier et al., 2017, Eshraghi, Kim et al., 2016, Lagrange, Benaoudia et al., 2018).

The transcriptional and post-transcriptional regulatory processes, controlling *Francisella* phagosomal escape and cytosolic multiplication, have been already well characterized (Celli & Zahrt, 2013, Charity, Blalock et al., 2009, Cuthbert, Ross et al., 2017, Meibom, Barel et al., 2009, Sjostedt, 2011, Ziveri, Barel et al., 2017). Notably, the crucial role of a 30-kb locus in *Francisella* virulence (**Figure supplement 1**), designated FPI for “Francisella pathogenicity island” (Broms, Sjostedt et al., 2010), has been extensively documented (Clemens, Lee et al., 2018). Of note, the FPI is duplicated three subspecies of *F. tularensis* (subsps *holarctica*, *mediasiatica*, and *tularensis*), but is present in a single copy in *F. novicida* and *F. philomiragia*

The FPI encodes a Type VI secretion sytem (T6SS) that is essential to promote bacterial phagosomal escape and access to the cytosolic replication niche (Lauriano, Barker et al., 2004, Meyer, Broms et al., 2015, Nano, Zhang et al., 2004). Whereas the transcriptional regulation of the FPI locus has been deeply characterized (Ramsey, Osborne et al., 2015), the molecular mechanisms triggering T6SS assembly and contraction remain largely unknown. Bioinformatic analyses have divided bacterial T6SS in three phylogenetically distinct subtypes (designated T6SSi-iii) (Russell, Wexler et al., 2014). Whereas most of the well-characterized T6SS belong to the T6SSi (including in *P. areruginosa* and *V. cholerae*), *Francisella* is currently the only bacterium to possess a T6SSii (Abby, Cury et al., 2016) known to be exclusively involved in the intracellular life cycle of the pathogen. Phagosomal escape occurs as early as 30 minutes post-infection (Pizarro-Cerda, Charbit et al., 2016). The FPI encodes, in addition to the T6SS components, a number of proteins of unknown functions some of which (such as IglF,PdpC or PdpD) have been proposed to be T6SS effectors (Eshraghi et al., 2016). Very recently, Mougous and co-workers elucidated the function of one substrate of the T6SS (Ledvina, Kelly et al., 2018). This effector (named OpiA for outside pathogenicity island protein A) is a phosphatidylinositol (PI) 3-kinase that acts on the Francisella-containing phagosomal membrane to promote bacterial escape into the cytoplasm.

Our current knowledge of the structure and functional assembly of the *Francisella* T6SS are mainly based on recent cryo-electron microscopy data (Clemens, Ge et al., 2015, Nazarov, Schneider et al., 2018) and structural homologies with other members of the T6SSi (Clemens et al., 2015, Nazarov et al., 2018) and references therein). It has been proposed that the tube comprised of IglC subunits (Hcp homologue) that form hexameric rings, fits within the cavity of the IglA/IglB sheath (TssB/C in *Pseudomonas*, VipA/B in *Vibrio*). Contraction of the sheath results in the ejection of the Hcp tube, together with effector proteins located within the tube and on top of the tip of the tube (Vettiger & Basler, 2016). Formation of the contracted *Francisella* T6SS can be mimicked in vitro by high K^+^ concentration (Clemens et al., 2015). Yet, the molecular relay for this environmental cue remains uncharacterized.

In order to understand the mechanisms underlying this KCl-induced T6SS production, we performed a global proteomic approach on KCl-induced and non-induced bacteria. This analysis revealed that the relative amounts of most FPI proteins were not increased upon KCl stimulation that suggested the possible implication of a post-translational control of T6SS polymerization. We hypothesized that protein phosphorylation might occur in *Francisella* and contribute to the dynamics of T6SS assembly-disassembly. By using state-of-the art mass spectrometry analyses, we indeed identified more than one hundred peptides bearing serine/threonine/or tyrosine phosphorylated residues in the proteome of *F. novicida.* One peptide, corresponding to a phosphorylation of IglB on tyrosine 139 (Tyr139), particularly attracted our attention. IglB is analogous to the TssC protein in canonical T6SS and forms with IglA (TssB homologue) the T6SS contractile sheath (Clemens et al., 2015). The data presented here reveal that phosphorylation of IglB is essential for the proper activity of the T6SS and could contribute to the fine control of T6SS dynamics.

## Results

We first monitored sheath formation upon KCl stimulation, using the procedure previously described (Clemens et al., 2015). Briefly, lysates of wild-type *F. novicida,* grown in the presence of 5% KCl, were layered onto 15%-55% sucrose gradients (**Figure 1A**). The fraction sedimenting in equilibrium at 55% sucrose and below (55% Optiprep, grey arrow) from KCl-induced cultures were further dialyzed and used for negative staining TEM imaging. As expected, rod-shaped particles were detected by transmission electron microscopy in this fraction that corresponded to contracted sheath-like structures. The different fractions of the gradient were next tested by western blotting with anti-IglA and anti-IglB antibodies. IglA and IglB were detected in all the fraction of the gradient in KCl-induced conditions (**Figure 1B**). In contrast, they were only detected in the fractions of the upper half of the gradient (in the 15% and in part of the 35% sucrose fractions) in the non KCl-induced conditions. We next decided to perform a whole cell proteome analysis of KCl-induced and non-induced bacterial cultures.

**Figure 1.**
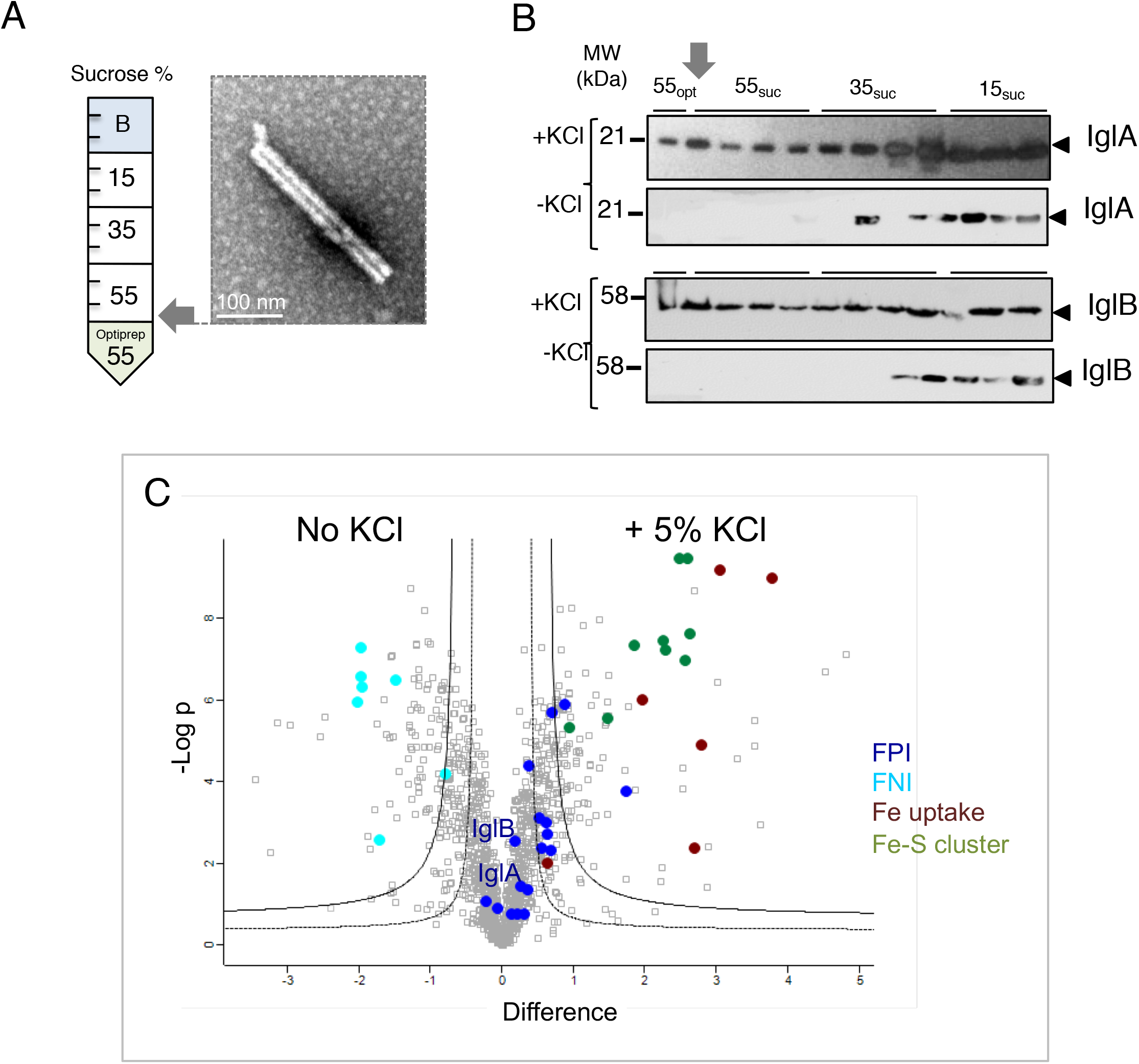
KCl stimulation of sheath formation. (**A**) Sucrose gradient. Lysates of bacteria grown in Schaedler K3 medium in the presence of KCl were laid on top of a discontinuous sucrose gradient 15%-55%/Optiprep 55%. Left panel, composition of the gradient; Right panel, transmission electron microscopy (TEM) of the “heavy” fraction (grey arrow, topping the Optiprep cushion). The assembled sheath-like structure, sedimenting at equilibrium to below 55% sucrose, showed rod-shaped particles of variable length (100 - 600 nm). (**B**) Western blotting analysis of the different fractions, using anti-Igl (A, B or C) antibodies. (**C**) The proteome of KCl-stimulated and non-stimulated *F. novicida.* Volcano plot representing the statistical comparison of the protein LFQ intensities of KCl-stimulation vs non-stimulated cells. Inner volcano was established using S0=0.5, FDR=0.01 (class B proteins) and the outer volcano using S0=0.5, FDR=0.001 (class A proteins). Proteins belonging to the FPI, the Fe uptale and Fe-S clusters are highlighted in color as indicated.

### Whole cell proteome analyses

Whole cells extracts from KCl-induced and non-induced cultures were analyzed in order to identify proteins whose intensity was altered by the KCl treatment. We could quantify 1,395 proteins that represent 80% of the predicted *Francicella* proteome (**Figure 1C**). Upon KCl stimulation, 113 proteins were up-regulated after KCl stimulation and 97 down-regulated, confirming that KCl stimulation strongly alters the proteome of the cell (**Table supplement 1**). Remarkably, the amounts of most of the FPI-encoded proteins (17 out the 18 proteins encoded by the FPI were identified) did not change significantly upon KCl stimulation, including notably the sheath and tube proteins IglA, IglB, IglC (blue dots in **Figure 1C**). The down-regulated proteins comprised seven proteins (FTN_0042 to FTN_0048) encoded by the “Francisella novicida island” (or FNI), a genomic island that shows some similarities with the FPI (Rigard, Broms et al., 2016), FTN_0042 and FTN_0043 corresponding to the orthologues of IglA and IglB, respectively). Oppositely, two sets of proteins emerged as up-regulated (**Figure supplement 2A**): proteins involved in Iron-sulfur (Fe/S) clusters biogenesis (FTN_0751 to FTN_0754, FTN_0850 to FTN_0853 and FTN1082) as well as most of the protein encoded by the *fig* (also designated *fsl*) operon (Kiss, Liu et al., 2008, Sullivan, Jeffery et al., 2006), involved in iron acquisition (FTN_1681, FTN_1682, FTN_1684 to FTN_1687) (**Figure supplement 2B**). On the basis of the known regulation of Fe/S clusters in other bacterial species (Mettert & Kiley, 2015, Roche, Aussel et al., 2013), these Fe/S cluster proteins could be regulated by FTN_0810, a predicted transcriptional regulator sharing modest sequence identity with the Fe/S regulator IscR.

### The phosphoproteome of *Francisella*

The fact that the amounts of T6SS proteins remained unchanged upon KCl induction, led up to hypothesize that a post-translational mechanism might be responsible for KCl-dependent sheath polymerization. We decided to test if protein phosphorylation could be involved in the process and carried out a global and site-specific phosphoproteomic analysis of *F. novicida* (strain U112), based on phosphopeptide enrichment and high-resolution LC-MS/MS analysis of whole cell lysates from KCl-induced and non-induced cultures (**Figure 2**). Overall, this analysis allowed the identification of 103 phosphopeptides, of which 78 were robustly quantified. The majority of the phosphosites (47 out of 78) were found on serine residues (S), while similar lower number of sites, 19 and 12, were found on threonine (T) and tyrosine (Y), respectively (**Figure 2A, B; Tables supplement 1**, **2**). These phosphosites corresponded to 59 proteins, of which 7 presented multiple phosphorylation sites. These proteins belonged to various functional categories and the most represented classes were carbohydrate metabolism, energy production and conversion and translation.

**Figure 2.**
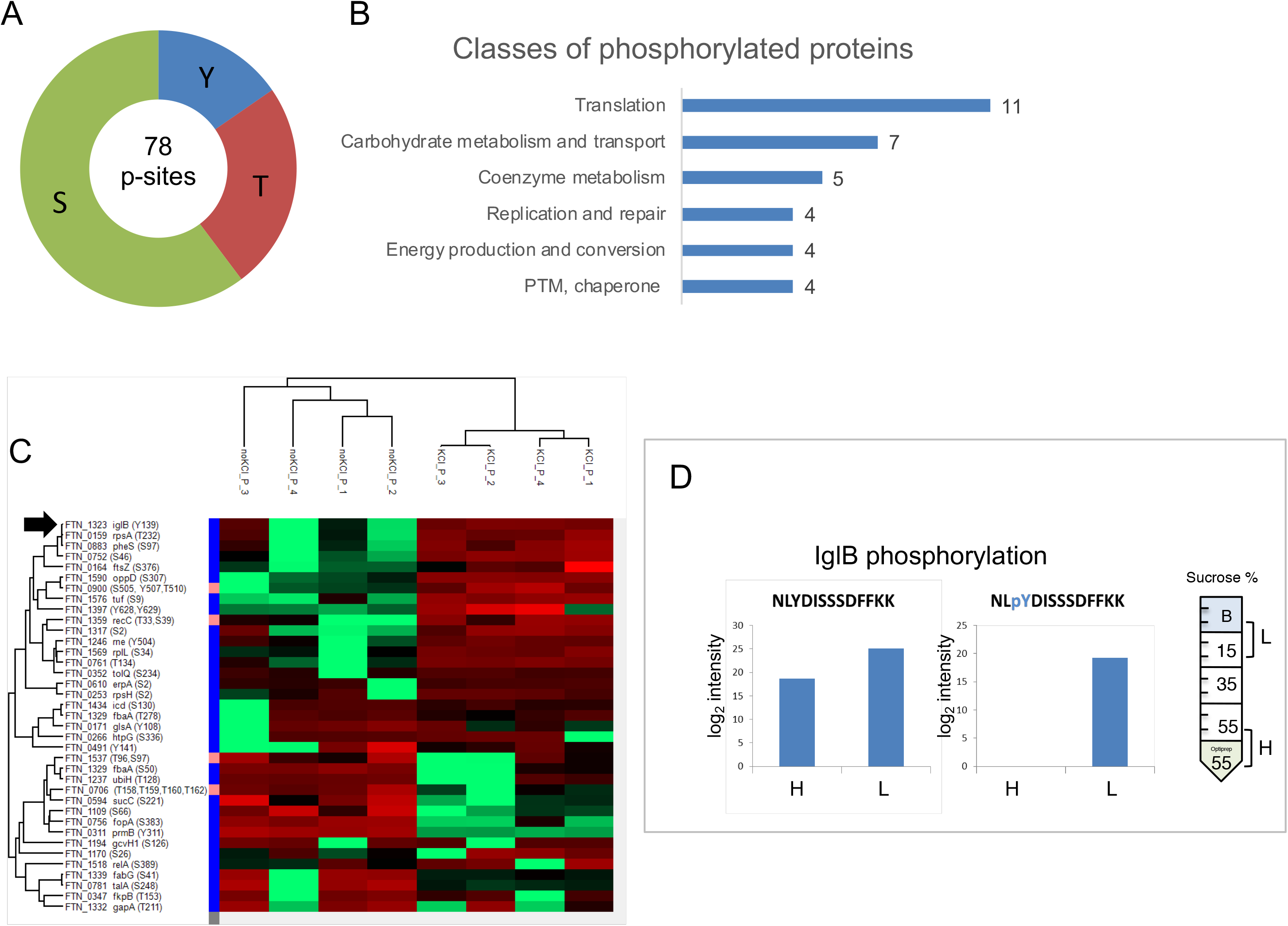
The phosphoproteome of *F. novicida*. (A) Distribution of phosphosites according to the modified amino acid (tyrosine,Y; threonine, T and serine, S). (B) Histograms of the most represented classes of protein. The values correspond to the number of proteins bearing phosphosites in each category. (C) Heat map of phosphoproteins altered upon KCl stimulation (Pink to right of the FTN numbers corresponds to proteins with multiple phosphorylation sites). (D) IglB phosphorylation distribution. Intensities (in log2) of the non phosphorylated (left) and phosphorylated (right) Tyr139-containing peptide in the “heavy” (H) and “light” (L) fractions. The signal of the phosphopeptide was only detectable after phosphoenrichment. nd, not detectable.

IglB was the only protein of the T6SS to be phosphorylated (**Figure 2C**). Furthermore, IglB which constitutes with IglA the T6SS sheath of *Francisella* (Broms et al., 2010, Clemens et al., 2015, Rigard et al., 2016) bore a unique phosphorylation site on Tyr139 (**Table supplement 3**). In order to determine if the phosphorylated forms of IglB was incorporated into the sheath upon its assembly, we further analyzed the fractions at the bottom of the gradient (sucrose 55%-Optiprep 55% density zone), containing the sheath-like particles) to the fractions at the top of the gradient (15-35% sucrose density zone, presumably containing non-polymerized forms of the IglA and IglB proteins (hereafter called “heavy fraction”, H or “light fraction”, L, respectively) (**Figure 2D**). The mass spectrometry analysis of the H and L fractions identified over 1,145 proteins (**Table supplement 4**) and confirmed the efficiency of the sucrose gradient separation. Unmodified peptides containing Tyr139 were detected in both H and L fractions, indicating that soluble IglB include both phosphorylated and non-phosphorylated forms of the protein. Importantly, the peptide bearing the phosphorylated tyrosine residue (pY139) of IglB was only detected in the L fraction of the gradient (**Table supplement 4**), suggesting that tyrosine phosphorylation might be unfavorable for sheath polymerization.

Fully supporting this notion, our 3D predictions (**Figure 3**) indicate that, upon addition of a phosphoryl moiety at this position, charge repulsion between the phosphate group and nearby aspartate residues should be unfavorable for the formation of stable hexameric rings and sheath contraction. Since the phosphate on Y139 is linked to the O^4^ position of the phenolic ring, the effects of Tyr phosphorylation may be exerted primarily through allosteric/electrostatic effects. Looking at the full helical reconstruction, one can see that Y139 is close to both D63 of IglA and D114 of IglB. These are from two different symmetry-related chains (**Figure 3**). Putting a phosphate on Y139 is thus likely to repel the aspartate residues and to disassemble the structure.

**Figure 3.**
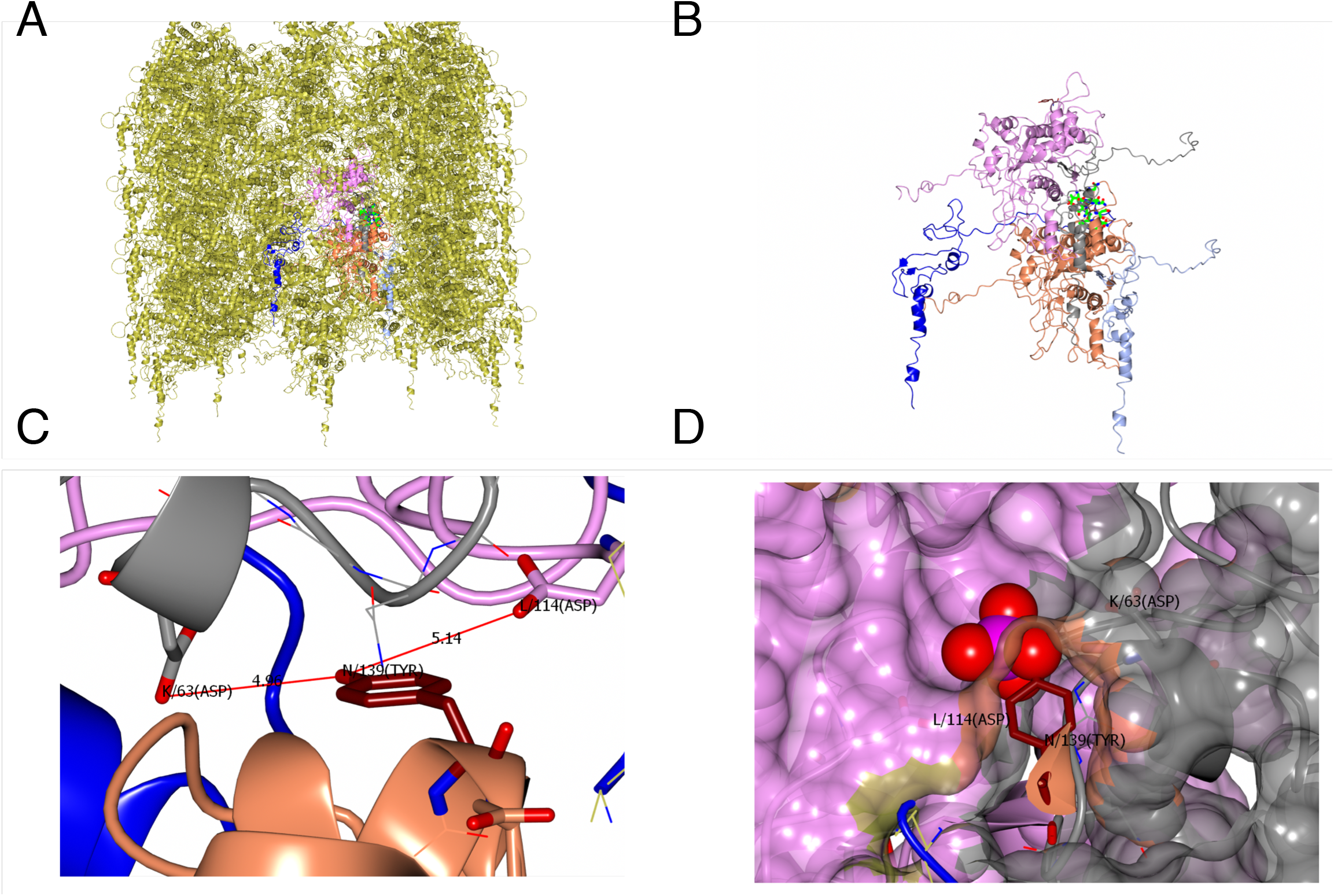
Predicted role of Y139 of IglB in sheath assembly. (**A**) Overview of the sheath that consists of 12 helical strands based on EM reconstruction at 3.7 Å. Each unit consists of a IglA/IglB dimer. The same colors were used in the further pictures. The chains near Tyr139 of IglB consist of 2 IglA/IglB dimers from the same strand (pink/grey and coral/sky blue). The N-terminal tail of an IglA (blue) from an adjacent strand wraps round to close to Y139 of this strand. (**B**) A closer view of just the selected chains. **(C)** Close up of the region of Tyr139. The hydroxyl is pointing into a pocket and not obviously making a Hydrogen bond. However, the carboxylates of Asp114 from IglB and Asp63 of IglA in the next dimer are around 5 Å away. At the resolution of the reconstruction side chain density is not well defined and these could in fact be close enough to H bond. **(D)** Surface of the assembly clipped to show Tyr139. There is enough space for a Phospho group on the Tyr. The two Asps will however be close to the phosphogroup and so there is likely to be charge repulsion impairing the sheath assembly or weakening the sheath stability when Y139 is phosphorylated.

### Sequence analyses

Comparative sequence analyses on 33 IglB proteins from 19 genomes of the *Francisella* genus were performed to assess the conservation of residue Tyr139 (**Figure supplement 3**). Sequences were highly conserved between 31 IglB proteins sharing 93 to 100% amino acid identity. In contrast, the proteins encoded by *FTN_0043* in the FNI (Rigard et al., 2016) and *F7308_1917* (Fsp_TX077308) exhibited only 48% amino acid identity with the 31 other IglB proteins whereas they both share 85% identity. Excluding these 2 outliers, among 506 sites, 458 were without polymorphism (91%), with Tyr139 conserved in all IglB sequences (**Figure supplement 3, upper part**). Since IglB is among the most conserved components of T6SS (Broms et al., 2010, Clemens et al., 2015), we were able to retrieve orthologues of IglB encoded outside of *Francisella* genus. Out of 535 non-redundant IglB orthologues (also named VipB, TssC or EvpB/VC_A0108), we found 90 sequences with a tyrosine in the vicinity of Tyr139 site outside of *Francisella* genus. In 82 sequences, the predicted location of the tyrosine residue was similar to that of Tyr139 (in a turn region between two helices) (**Figure supplement, lower part**).

We hereafter thoroughly evaluated the importance of Y139 phosphorylation on T6SS functionality and *Francisella* pathogenicity. For this, we constructed two IglB variants in which Tyr139 was substituted either by an alanine (Ala) or by the non-phosphorylatable aromatic amino acid analogue of tyrosine, phenylalanine (Phe). These IglB mutated proteins (designated Y139A and Y139F, respectively) were expressed *in trans* in a ∆*iglB* mutant of *F. novicida* carrying a chromosomal deletion of the entire *iglB* gene, (see Materials and Methods). A *F. novicida* mutant with a deletion of the entire *FPI*, unable to escape from phagosomes and hence to grow in macrophages (Weiss, Brotcke et al., 2007), was used as a negative control.

### A role of IglB phosphorylation in sheath formation

IglB was detected by Western blotting on whole cell lysates in wild-type and complemented mutated strains (Y139A and Y139F) but not in the ∆*iglB* and ∆*FPI* strains (**Figure 4A**). IglA was also detected in wild-type and complemented mutated strains (Y139A and Y139F), but in lower amounts in the ∆*iglB* strain and not in the ∆*FPI* mutant. This observation is in agreement with earlier observations (Broms, Lavander et al., 2009) that suggested that the presence of IglB increases the stability of IglA.

**Figure 4.**
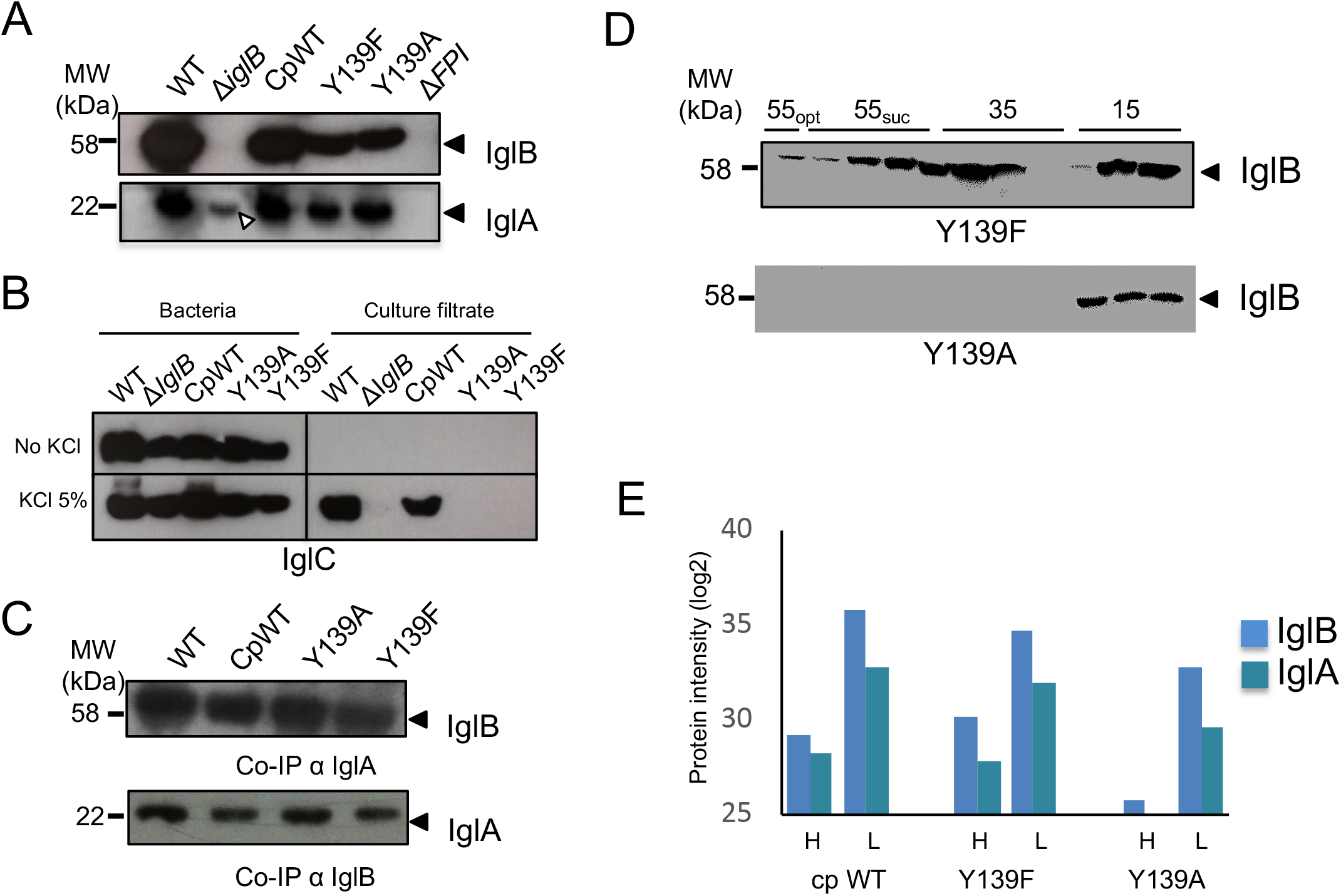
Immunodetection of IglA, IglB and IglC in wild-type and IglB mutant strains. (**A**) Western Blots (WB) of whole cell protein extracts. Upper gel: WB with anti-IglB; lower gel: WB with anti-IglA. (**B**) Western Blots (WB) with anti-IglC on bacteria / culture filtrates. Bacteria were grown in Schaedler-K3 supplemented or not with 5% KCl until late log phase and then harvested by centrifugation. Culture supernatants were collected after filtration on 0.2 µM Millipore filters and concentrated using Amicon 3 kDa membranes. The equivalent of 200 µg of total protein were loaded onto each well. Upper gel, bacterial cultures grown without KCl; lower gel, bacterial cultures supplemented with 5% KCl. (**C**) Co-immunoprecipitations (Co-IP). Upper gel: IP-anti-IglA, followed by WB with anti-IglB; lower gel: IP-anti-IglB, followed by WB with anti-IglA. (**D**) WBs of sucrose gradient fractions of the Y139F and Y139A mutant strains. **(E)** Distribution of the intensities (trasformed in log 2) of the proteins IglA and IglB in the Heavy (H) and light (L) sucrose fraction, in cp WT (∆*iglB* complemented with WT IglB), Y139F and Y139A (∆*iglB* complemented with IglB Y139F and IglB Y139A, respectively).

A classical measure of basal T6SS function is the export of an Hcp-related protein. Therefore, we examined IglB-dependent IglC export in the different *F. novicida* mutant strains by Western-blot on bacteria grown in the presence -or absence-of 5% KCl. In agreement with previous reports, IglC was detected in the culture supernatant in the presence, but not in the absence, of KCl, in the wild-type and *iglB*_WT_ complemented strains. In contrast, deletion of *iglB* as well as the two single amino acid substitutions Y139A and Y139F abolished the secretion of IglC into the culture medium (**Figure 4B**). KCl treatment did not modify the amounts of IglA, IglB and IglC proteins detected by western blotting on whole cells (bacterial pellet fractions).

To check the impact of the two amino acid substitutions on IglA/IglB heterodimer formation, we performed co-immunoprecipitation assays (**Figure 4C**). We used either anti-IglA to precipitate the complex, followed by western blotting with anti-IglB (upper panel) or anti-IglB, followed by western blotting with anti-IglA (lower panel). In both conditions, the two IglB mutants were still able to interact with IglA to form IglA/IglB heterodimers.

We next performed sucrose gradient fractionation (as described above), followed by western blotting with anti-IglB on the Y139F and Y139A mutant strains. The Y139F IglB mutant protein was detected in both the L and H fractions of the gradient (**Figure 4D**), indicating that some sheath polymerization still occurred in the Y139F mutant. In contrast, the Y139A mutant protein was detected in the upper fractions of the gradient but not in the lower fractions (corresponding to the contracted sheath). Fully supporting the western-blot data, proteomic analysis of the distribution of IglA and IglB in the Heavy (H) and light (L) sucrose fraction revealed that their distribution was similar in the strains expressing either WT IglB and Y139F IglB with, in both case, a slightly higher proportion of IglA and IglB in the L fraction compared to the H fraction. In sharp contrast, with the Y139A mutant, IglB was absent from the H fraction and IglA also almost exclusively found in the L fraction (**Figure 4E)**.

Altogether, these data comforted the infection assays described above and suggested that the non-phosphorylatable Y139F mutant could still assemble into not fully functional sheath-like structures.

### Critical role of residue Y139 in *Francisella* pathogenesis

The mutations Y139A and Y139F had no effect on bacterial growth in broth (**Figure supplement 4**). The ability of wild-type *F. novicida* (WT), ∆*iglB* and complemented strains (CpWT, Y139A and Y139F) to survive and multiply in murine macrophage-like J774.1 cells was monitored over a 24 h-period (**Figure 5A**). Confirming earlier reports (Spidlova & Stulik, 2017), intracellular multiplication of the Δ*iglB* mutant was essentially abolished and comparable to that of the ∆*FPI* mutant. Remarkably, the single amino acid substitution of Tyr139 by Ala (Y139A) also abolished intracellular bacterial multiplication. In contrast, when Tyr139 was substituted by Phe (Y139F), a severe intracellular growth defect was recorded until 10 h after infection (almost fifty-fold less bacterial counts) but at 24 h, the Y139F mutant strain had multiplied and showed only a ten-fold reduction of bacterial counts compared to wild-type, suggesting that a late phagosomal escape had occurred therefore promoting cytosolic bacteria multiplication.

**Figure 5.**
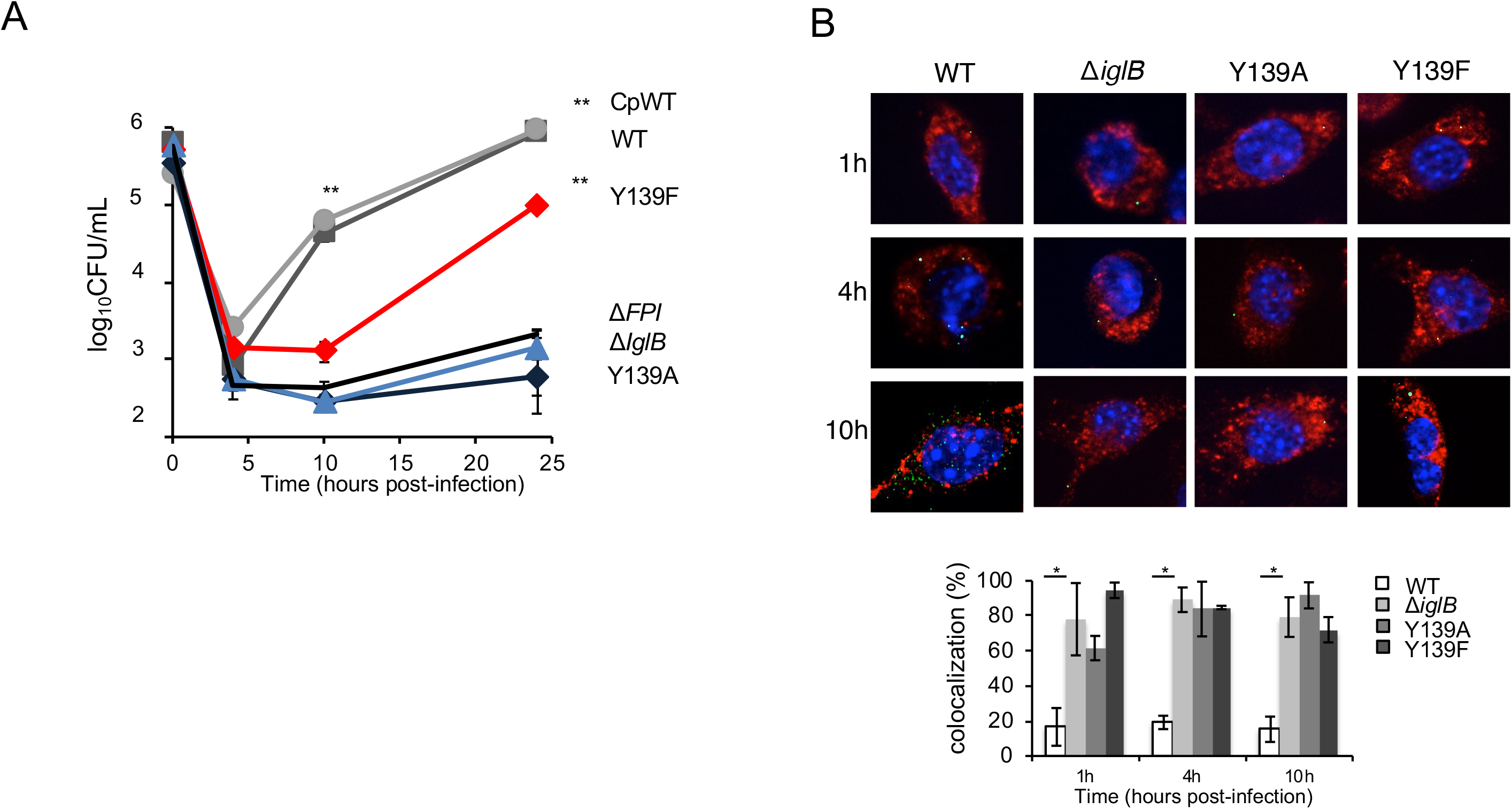
Y139Aand Y139F IglB mutants show impaired intracellular survival and attenuated virulence. (**A**) Intracellular bacterial multiplication of wild-type *F. novicida* (WT, Grey squares), isogenic ∆*iglB* mutant (Δ*iglB*, blue triangles) and complemented Δ*iglB* strain (Cp-WT, grey circles; Y139A, blue square; Y139F, red squares) and the Δ*FPI* control (black lines), was monitored during 24 h in J774A.1 macrophage cells. The values recorded for Y139F, at 24 h, were significantly different from those of ∆*FPI*, ∆*iglB* or Y139A. In both cell types, the values recorded for WT or CpWT, at 10 h and 24 h, were significantly different from those of ∆*FPI*, ∆*iglB,* Y139A or Y139F. **p<0.001 (determined by student’s t-test). (**B**) Confocal microscopy of intracellular bacteria with J774-1 infected with wild-type *F. novicida* (WT), Δ*iglB* or complemented strains and their co-localization with the phagosomal marker LAMP1 observed at 1 h, 4 h and 10 h. Upper par: J774.1 were stained for *F. novicida* (green), LAMP-1 (red) and host DNA (blue, DAPI stained). Lower part: analysis was performed with ImageJ software. *p<0.01 (determined by student’s t-test).

As expected, functional complementation (*i.e.,* introduction of a plasmid-born wild-type *iglB* allele into the ∆*iglB* mutant strain (CpWT) restored wild-type growth.

We compared the ability of the ∆*iglB* mutants to escape from the phagosomal compartment to that of the wild-type strain by monitoring co-localization of intracellular bacteria with LAMP-1 in J774-1 macrophages (**Figure 5B**), using specific antibodies and automated quantification. The frequency of bacteria co-localizing with LAMP-1 at all 3 time points was elevated (between 60% and 90%), with ∆*iglB* and the two complemented strains expressing mutated *iglB* alleles (designated Y139A and Y139F; respectively). In contrast, co-localization of the wild-type strain with LAMP-1 was below 20% after 1 h and remained in the same range throughout the infection, suggesting that the ∆*iglB* mutant as well as the two strains expressing mutated *iglB* alleles, are still trapped in the phagosomal compartment after 10 h, whereas the wild-type strain has already escaped into the cytosol after 1 h.

However, supporting the kinetics data, transmission electron microscopy confirmed that a clear cytosolic multiplication was visible at 24 h with the Y139F mutant but not with the Y139A mutant (**Figure 6A**). We therefore next used time-lapse video microscopy to visualize in real time multiplication of the Y139F mutant in J774-1 macrophages. For this, we inserted a GFP-encoding gene into plasmids pKK214::pGro*iglB* and pKK214::pGro*iglB*Y139F (yielding recombinant plasmids pKK214::pGro*iglB*-pGro*gfp* and pKK214::pGro*iglB*Y139F-pGro*gfp,* respectively). *F. novicida* ∆*iglB* was transformed with these GFP-expressing recombinant plasmids. We generated J774-1 cells with red nuclei by transduction with NucLight Red Lentivirus to facilitate cell recognition and counting (see Materials and Methods). J774-1 cells with red nuclei were infected by GFP-expressing bacteria at an MOI of 1,000. Infection was followed over a 24h-period, using a fully automated microscope Incucyte^®^ 531 S3 (Essen BioScience). Images were taken every 20 minutes with the 20X objective during 24 h, starting 1 h after infection (**Figure 6B** snapshots; **Videos supplement 1, 2**). Under these conditions, an active intracellular multiplication was observed with both strains. In the conditions used, we observed only a minor delay in bacterial multiplication was observed with the strain complemented with the mutated *iglB* allele Y139F compared to the strain complemented with the wild-type allele, In contrast to the kinetics and LAMP co-localization data that suggested a retarded phagosomal escape of the Y139F mutant. Hence, it is likely that the Y139F substitution allows the production of sufficient assembled T6SS to promote bacterial phagosomal escape and cytosolic multiplication in these cells.

**Figure 6.**
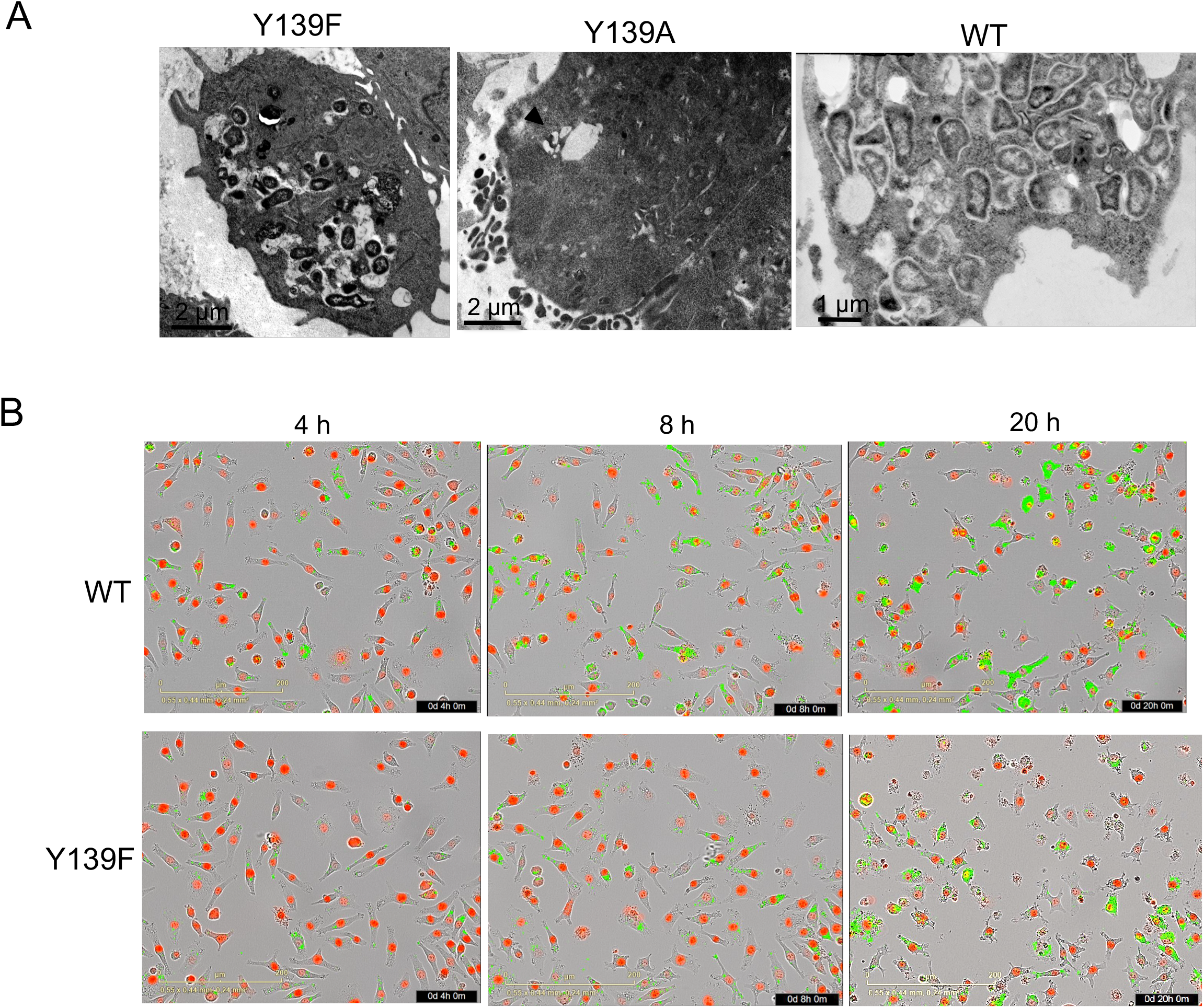
Electron microscopy and time lapse video microscopy. (**A**) Transmission electron microscopy of J774-1 cells infected with Y139F, Y139A or WT *F. novicida* at 24 h. Black arrowheads (in cells infected with Y139A) point to single bacteria. (**B**) Time lapse video microscopy. J774-1 cells expressing a nuclear restricted Red Fluorescent Protein were infected with GFP-expressing WT and Y139 IglB proteins. Representative images were taken at 4,h, 8 h and 20 h (see Video S1 and S2). Red: cell nuclei. Green: GFP expressing bacteria.

**Figure 7.**
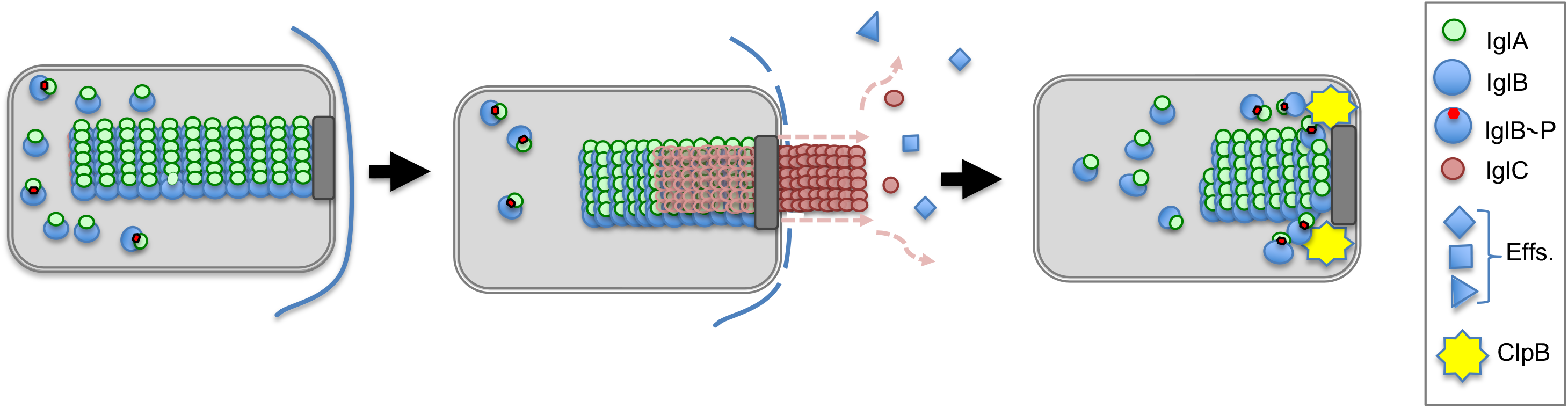
Proposed model of phosphorylation-dependent TSS6 disassembly. Upon bacterial entry into the phagosomal compartment, non-phosphorylated IglB form stable heterodimers with IglA and sheath polymerization initiates at the bacterial pole. Within 30 min, T6SS sheath contraction triggers the secretion of effectors and the rupture of the phagosomal membrane. Immediately after, the T6SS sheath disassembles, thanks to the concerted action of the ClpB unfoldase and of the phosphorylation of IglB residue Y139.

Finally, we replaced Tyr139 by two phosphomimetics *i.e.* the negatively charged amino acids Aspartate and Glutamate and evaluated the impact of these amino acid substitutions (Y139D and Y139E) on intracellular multiplication in J774-1 cells (**Figure supplement 6A**). The two mutants led to severe defects in intracellular multiplication and, at all time points tested, the replication defect was comparable to that of the Y139A or ∆*FPI* mutant. We next evaluated the capacity of the Y139D and Y139E mutants to form sheath-like structure by using the sucrose gradient assay. In the two mutants, the IglB protein was exclusively detected by Western blot in the upper fractions of the sucrose gradient (**Figure supplement 6B**). These data strongly suggest that residues possibly mimicking the phosphorylated state of Y139 also prevented sheath-like assembly.

Of note, in primary bone marrow-derived macrophages (BMMs), growth of the ∆*iglB* deletion mutant as well as that of the two Y139A or Y139F was similar to that of a ∆*FPI* mutant (**Figure supplement 5A**) at all time points tested. Functional complementation restored normal intracellular bacterial replication (in CpWT). The ability of the ∆*iglB* mutants to induce phagosomal membrane rupture in BMMs was further tested by using a CCF4 assay, essentially as previously described (Ramond, Gesbert et al., 2015, Rigard et al., 2016). Wild-type *F. novicida* and CpWT strains showed similar amount of cleaved CCF4 at the three time-points tested (1 h, 3 h and 6 h) whereas there was no (or marginal) loss of FRET signal with ∆*iglB* (**Figure supplement 5B**), Y139A, Y139F, and the two negative control strains (∆*FPI* and ∆*bla*), indicating that the Y139A, Y139F mutant strains remained stuck in intact phagosomes during the time-period tested in BMMS. Hence in primary macrophages, a tyrosine residue is required for an optimal T6SS assembly likely due to their high bactericidal activity. Moreover, in the mouse model of tularemia (see Materials and Methods for details), the two mutants Y139A and Y139F also showed an almost complete loss of virulence (**Fig S5C)** that confirmed the critical role of tyrosine residue 139 of IglB in Francisella pathogenicity.

### Searching for putative kinases

Our careful inspection of the *Francisella* genomes did not identify any gene encoding for a putative serine/threonine- or tyrosine-kinase. We therefore searched for proteins containing potential tyrosine kinase domains. Using the PFAM database (https://pfam.xfam.org/), we could identify three candidates proteins (FTN_1082, FTN_0459 and FTN_1433). FTN_1082 is predicted to belong to the uncharacterized ACR,YdiU motif/UPF0061 family (PFAM database PF02696). This protein family notably comprises the SELO proteins (for selenoproteins) that have been proposed to constitute a novel ubiquitous kinase-like family of proteins. Recent bioinformatic analyses have identified FTN_1082 as a potential SELO family-member (Dudkiewicz, Szczepinska et al., 2012). FTN_0459, is predicted to encode a UbiB homologue, The UbiB protein family is also a widespread family of kinase-like present in the three kingdoms of life, often required for the biosynthesis of isoprenoid lipids. In *Escherichia coli,* UbiB has been shown to be required for the aerobic biosynthesis of the redox-active lipid ubiquinone, coenzyme Q (Aussel, Pierrel et al., 2014). It has been assumed but not formally established that UbiB family members were protein kinases. However, very recently it was demonstrated that a UbiB family member, the mitochondrial protein ADCK3, adopted a PKL fold and bound adenine nucleotides in a divalent cation-dependent manner strongly supporting the hypothesis that UbiB proteins are bona fide kinases (Stefely & Pagliarini, 2017). Finally, FTN_1433, is a protein bearing a G-rich domain identified on putative tyrosine kinase (PFAM database PF02706). This family notably includes proteins involved in lipopolysaccaride biosynthesis (Wzz proteins).

We anticipated that if one of these genes was involved in IglB phosphorylation, its inactivation would impair phagosomal escape and hence cytosolic bacterial multiplication. We therefore monitored intracellular bacterial multiplication of the three corresponding transposon insertion mutants of *F. novicida* (Gallagher, Ramage et al., 2007) after 10 hours in J774-1 macrophages (**Figure supplement 7**). The three mutants showed wild-type intracellular multiplication in this assay, suggesting that either these proteins work synergistically or that there exists another yet unidentified enzyme with a moonlight tyrosine kinase activity.

## Discussion

We show here that the unique phosphorylated residue of the sheath component IglB plays a critical role in T6SS functional assembly, bacterial phagosomal escape and virulence. This post-translational modification of IglB might contributes to the fine tuning of T6SS disassembly.

### Pivotal role of residue 139 in sheath assembly

The assembly of the T6SS-H1 of *Pseudomonas aeruginosa* has been shown to be spatially regulated by a post-translational mechanism coined « the Threonine Protein Phosphorylation » (or TPP) Pathway. During this process, involved in the defense of bacteria to a neighboring attack, a membrane-bound threonine kinase phosphorylates a forkhead-associated domain-containing Protein (FhA or TagH), which promotes the assembly of an active conformation of the T6SS at the site of the attack (Cianfanelli, Monlezun et al., 2016, Mougous, Gifford et al., 2007). More recently, the offensive anti-bacterial T6SS of *Serratia marcescens* has also been shown be controlled by the opposing actions of the TPP pathway and TagF on assembly of the core machinery (Ostrowski, Cianfanelli et al., 2018). The TPP represents the only currently identified post-translational mechanism regulating T6SS biogenesis.

In bacteria such as *Escherichia coli, Vibrio cholerae* or *Pseudomonas aeruginosa*, the sheath contracts spontaneously during the repeated rounds of ultraspeed purification required for their isolation from cell cultures (Nguyen, Douzi et al., 2018). In *Francisella*, T6SS contraction can be triggered in culture either by placement under coverslips or by KCl stimultation (Clemens et al., 2015). Our mass spectrometry analyses revealed that T6SS proteins were not upregulated upon KCl treatment and that IglB was the only T6SS protein to contain a phosphorylated residue (Tyr139). Importantly, the phosphorylated form of IglB was not associated with the contracted sheath (**Figure 2**) that suggested that the phosphorylation status of IglB residue Tyr139 could participate to T6SS dissociation.

Formation of the *Francisella* T6SS is believed to occur in the phagosome before cytosolic escape of the bacterium and the T6SS seems to be dispensable for cytosolic multiplication. Nevertheless, some T6SS elements are likely to be required during the cytosolic stage of the infectious cycle. Indeed, most FPI genes are induced during late intra-macrophage growth (Wehrly, Chong et al., 2009) and IglC has been shown to be required for intracellular growth of *F. novicida* that are microinjected directly into the cytosol of HeLa cells (Meyer et al., 2015). IglC secretion in *F. novicida* depends on the T6SS core components IglA and IglB (Ludu, de Bruin et al., 2008). Of note, truncated IglA proteins still able to form IglA/IglB-heterodimers have been reported to have lost their IglC secretory function (Clemens et al., 2015) that suggests that heterodimer formation is not sufficient to generate a functional sheath.

The drastic impact of both Y139A and Y139F substitutions on bacterial virulence (**Figure supplement 5**) demonstrated that tyrosine residue 139 of IglB plays a critical role in the functional assembly of the T6SS. Remarkably, the Y139F mutant, which corresponds to the substitution of a tyrosine residue by its non-phosphorylatable aromatic analog phenylalanine, could still assemble unstable sheath-like structure (**Figure 4D, E**) and multiply in J774-1 macrophages (**Figure 5A; Figure 6A, B; video supplement 2**) and. At this stage, the inability of the Y139F mutant to form stable T6SS might be explained by the fact that it lacks the OH moiety on the C4 of the phenol ring. Indeed, several reports support the notion that the OH moiety of tyrosine residues may participate to protein functionality and stability. For example it has been shown that intramolecular hydrogen may contribute significantly to conformational protein stability (Bolen & Rose, 2008). Since the OH moiety is capable of forming such intra- or intermolecular hydrogen bonds with the carboxy oxygen of adjacent amino acid residues, when a Tyr-Phe substitution is made in a protein, it is likely to impact its folding or stability (Pace, Horn et al., 2001). More recently, (Baroni, Pandini et al., 2012) also demontrated that a single tyrosine Hydroxyl Group controled the specificity of *Plasmodium falciparum* ferredoxin-NADP+ reductase. Although inspection of the 3.7 Å EM reconstruction of the *F. novicida* contracted sheath (PDB 3j9o) does not show any atoms close enough to form a hydrogen bond with the hydroxyl of Tyr139, the carboxylates of Asp114 from IglB, and Asp63 of IglA in the next dimer, are around 5 Å away and these may approach close enough to Tyr139 OH to form hydrogen bond interactions stabilizing the assembly.

3D modeling also indicated that there is space for a phosphate group on this atom as the OH is pointing into a pocket in the structure (**Figure 3**). However, the negatively charged phosphate would induce charge repulsion with the nearby Asps from the next dimer most likely preventing assembly of the strands when Tyr139 is phosphorylated. Hence, the charge repulsion provoked by addition of the phosphogroup (when Y139 is phosphorylated) is likely to weaken the stability of the contracted sheath. Altogether these data suggest that the phosphorylation status of IglB could modulate T6SS assembly/disassembly process. Supporting the notion that phosphorylation of Tyr139 impairs sheath contraction, replacement of this residue by a phosphomimetic (Y139D or Y139E) led to a severe intracellular multiplication defect and prevented sheath-like structure formation.

### IglB phosphorylation, a new partner of T6SS dissociation

Basler and co-workers have recently shown that *F. novicida* assembled its T6SS sheath on the bacterial poles (Brodmann et al., 2017) and that the sheath cycled through assembly, contraction and disassembly, similarly to what was previously described for other canonical T6SSs. The FPI does not encode any homologue of the unfoldase ClpV, present in all the canonical T6SS, but instead uses the general chaperone ClpB for the recycling of the contracted sheaths (Brodmann et al., 2017). ClpB specifically localizes with the contracted sheaths and is responsible for their disassembly within phagosomes of infected macrophages. We found that the amounts of ClpB were not (or marginally) up-regulated upon KCl induction (*i.e.* only 1.3-fold more ClpB in KCl-induced conditions compared to non-induced conditions). ClpB was also mainly associated with the soluble fraction of the sucrose gradient of KCl-induced bacteria (**Table supplement 1**). The kinase activity, leading to IglB phosphorylation might assemble, like ClpB, at the pole of the bacterium only when the sheath is contracted and assist sheath dissociation.

In spite of our repeated attempts, we were unable to identify a candidate tyrosine kinase. This suggests that a yet unidentified enzyme with either a non-canonical tyrosine kinase activity or a protein predicted to be involved in a different enzymatic reaction with a « moonlight » tyrosine kinase activity, is responsible for IglB phosphorylation. If so, its identification might prove extremely difficult and would require extensive future work. It is also possible that a single kinase mutant might have no or very limited effect on phagosomal disruption since sheath polymerization would occur and be sufficient to promote phagosomal membrane disruption. Indeed, it has been shown (Meyer et al., 2015) that only a minor fraction of functional T6SS is sufficient to promote phagosomal escape and, once in the cytosol, FPI mutants can multiply like wild-type bacteria.

Our whole cell proteomic analysis revealed that the amounts of more than two hundred proteins were altered upon KCl stimulation (**Figure 1**). Notably, the down-regulated proteins comprized seven proteins (FTN_0042 - FTN_0048) out of the 13 proteins potentially encoded by the *Francisella novicida* island (FNI, FTN_0042 - FTN_0054) (Rigard et al., 2016). Although inactivation of the whole FNI locus had no effect on *F. novivcida* virulence (Rigard et al., 2016), a possible contribution of the orthologues of IglA and IglB (FTN_0042 and FTN_0043) to the sheath assembly-disassembly process cannot be excluded. The proteins involved in Iron-sulfur (Fe/S) clusters biogenesis or in iron acquisition, that were upregulated upon KCL stimulation might also contribute to T6SS biogenesis.

In conclusion, we have identified in the present work a critical residue of IglB whose phosphorylation may represent a new player involved in the fine-tuning of the T6SS dynamics of *Francisella*. Such post-translational sheath modifications might exist in other bacteria that need to be explored.

## Materials and Methods

### Ethics Statement

All experimental procedures involving animals were conducted in accordance with guidelines established by the French and European regulations for the care and use of laboratory animals (Decree 87–848, 2001–464, 2001–486 and 2001–131 and European Directive 2010/63/UE) and approved by the INSERM Ethics Committee (Authorization Number: 75-906).

### Strains and culture conditions

All strains used in this study are derived from *F. tularensis* subsp. *novicida* U112 (*F. novicida*) as described in **Table supplement 5**. Strains were grown at 37°C on pre-made chocolate agar PolyViteX plates (BioMerieux), Schaedler K3 or Chemically Defined Medium (CDM). The CDM used for *F. novicida* corresponds to standard CDM (Chamberlain, 1965) without threonine and valine (Gesbert, Ramond et al., 2014). For growth condition determination, bacterial strains were inoculated in the appropriate medium at an initial OD_600_ of 0.05 from an overnight culture in Schaedler K3 medium.

### Construction of a ∆ iglB *deletion mutant*

We inactivated the gene *iglB* in *F. novicida* (*FTN_1323*) by allelic replacement, resulting in the deletion of the entire gene (4 first codons and 7 last codons were conserved). We constructed a recombinant PCR product containing the upstream region of the gene *iglB* (*iglB*-UP), a kanamycin resistance cassette (*nptII* gene fused with *pGro* promoter) and the downstream region of the gene *iglB* (*iglB*-DN) by overlap PCR. Primers *iglB* upstream FW (p1) and *iglB* upstream (spl_K7) RV (p2) amplified the 689 bp region upstream of position +3 of the *iglB* coding sequence (*iglB*-UP), primers *pGro* FW (p3) and *nptII* RV (p4) amplified the 1091 bp kanamycin resistance cassette (*nptII* gene fused with *pGro* promoter); and primers *iglB* downstream (spl_K7) FW (p5) and *iglB* downstream RV (p6) amplified the 622 bp region downstream of the position +1520 of the *iglB* gene coding sequence (*iglB*-DN). Primers p2 and p5 have an overlapping sequence of 12 nucleotides with primers p3 and p4 respectively resulting in fusion of *iglB*-UP and *iglB-*DN with the cassette after cross-over PCR (**Table S6**). All single fragment PCR reactions were realized using Phusion High-Fidelity DNA Polymerase (ThermoScientific) and PCR products were purified using NucleoSpin^®^ Gel and PCR Clean-up kit (Macherey-Nagel). Overlap PCRs were carried out using 100 ng of each purified PCR products and the resulting fragment of interest was purified from agarose gel. This fragment was then directly used to transform wild type *F. novicida* by chemical transformation (Gesbert, Ramond et al., 2015). Recombinant bacteria were isolated on Chocolate agar plates containing kanamycin (10 μg mL^−1^). The mutant strains were checked for loss of the wild type gene by PCR product direct sequencing (GATC-biotech) using appropriate primers.

### Functional complementation

The ∆*iglB* mutant strain was transformed with the recombinant pKK214-derived plasmids (pKK-*iglBY139Acp*, pKK-*iglBY139Fcp*, pKK-*iglBY139Dcp* and pKK-*iglBY139Ecp*) described below (**Table supplement 5**). Primers pGroFW and pGro RV amplified the 328 bp of the *pGro* promoter; primers *iglB* FW/ and *iglB*[PstI] RV amplified the 1,534 bp *iglB* gene from U112. PCR products were purified and SmaI (*pGro* promoter) or PstI (*iglB*) digested in presence of FastAP Thermosensitive Alkaline Phosphatase (ThermoScientific) to avoid self-ligation. Mixtures of *pGro* promoter and fragments of the genes of interest were then incubated with T4 DNA Ligase (New England Biolabs) to allow blunt end ligation and fragments were then cloned in pKK214 vector after SmaI/PstI double digest and transformed in *E. coli* TOP10. Recombinant plasmid pKK-*iglBcp* was purified and directly used for chemical transformation in *F. novicida* ∆*iglB* (Gesbert et al., 2015). Recombinant colonies were selected on Chocolate agar plates containing tetracycline (5 μg mL^−1^) and kanamycin (10 μg mL^−1^).

For site-directed mutagenesis of *iglB* gene, we used plasmid pKK-*iglBcp.* The recombinant plasmids pKK-*iglBY139Acp*, pKK-*iglBY139Fcp*, pKK-*iglBY139Dcp* and pKK-*iglBY139Ecp* were constructed using the primer pairs iglB (Y/A, Y/F,Y/D, or Y/E) FW and iglB RV (**Table supplement 6**). The three first bases of primers iglB FW were modified to change Tyr139 to Ala, Phe, Asp or Glu PCR products were purified, phosphorylated by the T4 Polynucleotide Kinase and then incubated with T4 DNA Ligase (New England Biolabs) and transformed in *E. coli* TOP10. Recombinant plasmids were purified and directly used for chemical transformation in *F. novicida* ∆*iglB*, as described previously (Gesbert et al., 2015). Recombinant colonies were selected on Chocolate agar plates containing tetracycline (5 μg mL^−1^) and kanamycin (10 μg mL^−1^).

### Gfp-expressing *F. novicida*

The ∆*iglB* mutant strain was transformed with the recombinant plasmids pKK-GFP-*iglBWT_cp_*and pKK-GFP-*iglBY139F_cp_,* derived from pKK214-GFP (Abd, Johansson et al., 2003). Primers pGro (KlionskyAbdelmohsen et al.)FW and *iglB*(Klionsky et al.) RV (**Table supplement 5**) amplified the 1,870 bp of the *pGro-iglB* from the pKK-*iglBWT_cp_* and pKK-*iglBY139Fcp*. PCR products were purified and digested with SmaI and the *pGro-iglB WT* and *pGro-iglB Y139F* fragments were then incubated with T4 DNA Ligase (New England Biolabs) to allow blunt end ligation and fragments were then cloned in the unique SmaI site of pKK214-(GFP) vector and transformed in *E. coli* TOP10. Recombinant plasmid pKK-(GFP)-*iglBWT_cp_* and pKK-(GFP)-*iglBY139F_cp_* were purified and directly used for chemical transformation in *F. novicida* ∆*iglB* (Gesbert et al., 2015). Recombinant colonies were selected on Chocolate agar plates containing tetracycline (5 μg mL^−1^) and kanamycin (10 μg mL^−1^).

### Immunodetection

Antibodies to IglA, IglB and IglC were obtained through the NIH Biodefense and Emerging Infections (BEI) Research Resources Repository, NIAID, NIH.

#### Immunoblotting analysis

Protein lysates for immunoblotting were prepared by using Laemmli sample buffer. Protein lysates corresponding to equal OD_600_ were loaded on 12% Bis/Tris gels (Invitrogen), and run in TGS buffer. Primary antibodies (anti-IglA or anti-IglB) were used at final dilution of 1:2,000. Secondary horseradish peroxidase (HRP)-conjugated goat anti-mouse antibody (Santa Cruz Biotechnology, CA, USA) or (HRP)-conjugated goat anti-rabbit antibody and the enhanced Chemiluminescence system (ECL) (Amersham Biosciences, Uppsala, Sweden) were used as previously described (Ziveri, Tros et al., 2017).

#### Co-immunoprecipitations

Wild-type *F. novicida* was grown to late exponential phase in Schaedler K3; collected by centrifugation and lysed with lysozyme and 1% TritonX-100 detergent in 20 mM Tris HCl, (pH 7.8) with 1 mM EDTA, protease inhibitor cocktail and benzonase nuclease. The lysate was centrifuged at 15,000 g for 15 min at 4°C to pellet bacterial debris. Monoclonal anti-IglB antibody was incubated 40 min with Dynabeads protein G (Invitrogen) and the complex was incubated at room temperature with the bacterial lysate for 1 hour. After washing, the complex was loaded on 12% Bis/Tris gels (Invitrogen), and run in TGS buffer.

### T6SS purification

T6SS were prepared essentially as described in (Clemens et al., 2015). Briefly, wild-type *F. novicida* was grown to late exponential phase in Schaedler K3 containing 5% KCl; pelleted by centrifugation and lysed with lysozyme and 1% TritonX-100 detergent in 20 mM Tris HCl, (pH 7.8) with 1 mM EDTA, protease inhibitor cocktail and benzonase nuclease. The lysate was centrifuged 3 times at 15,000 g for 30 min at 4°C to pellet bacterial debris, and the supernatant was layered onto a 10%–55% sucrose gradient overlying a 55% Optiprep cushion and centrifuged at 100,000 g for 18 hr. Fractions were collected and examined by TEM negative staining using 2% uranyl acetate. The sheath-like structures sedimented to just below the 55% sucrose/Optiprep interface.

### Phosphoproteomic analyses

#### Reagents and chemicals

For protein digestion, dithiothreitol, iodoacetamide and ammonium bicarbonate were purchased from Sigma-Aldrich (St Louis, MO, USA). For phosphopeptide enrichment and LC-MS/MS analysis, trifluoroacetic acid (TFA), formic acid, acetonitrile and HPLC-grade water were purchased from Fisher Scientific (Pittsburgh, PA, USA) at the highest purity grade.

#### Protein digestion

For proteomic analysis, *F. novicida* was analysed in three independent biological replicates. Protein concentration was determined by DC assay (Biorad, CA, USA) according to the manufacturer’s instructions. An estimated 1.2 mg of proteins for each biological replicates were digested following a FASP protocol (Lipecka, Chhuon et al., 2016) slightly modified. Briefly, proteins were reduced using 100 mM dithiothreitol in 50 mM ammonium bicarbonate for 1h at 60°C. Proteins were then split into four samples of 300 µg and applied on 30 kDa MWCO centrifugal filter units (Microcon, Millipore, Germany, Cat No MRCF0R030). Samples were mixed with 200 µL of 8M urea in 50 mM ammonium bicarbonate (UA buffer) and centrifuged for 20 min at 15,000 × g. Filters were washed with 200 µL of UA buffer. Proteins were alkylated for 30min by incubation in the dark at room temperature with 100 µL of 50 mM iodoacetamide in UA buffer. Filters were then washed twice with 100 µL of UA buffer (15,000 × g for 15 min) followed by two washes with 100 µL of 50 mM ammonium bicarbonate (15,000 × g for 10 min). Finally, sequencing grade modified trypsin (Promega, WI, USA) was added to digest the proteins in 1:50 ratio for 16 h at 37°C. Peptides were collected by centrifugation at 15,000 × g for 10min followed by one wash with 50mM ammonium bicarbonate and vacuum dried.

#### Phosphopeptides enrichment by titanium dioxide (TiO_2_) and phosphopeptides purification by graphite carbon (GC)

Phosphopeptide enrichment was carried out using a Titansphere TiO_2_ Spin tip (3 mg/200 μL, Titansphere PHOS-TiO, GL Sciences Inc, Japan) an estimated 1.2 mg of digested proteins for each biological replicate. Briefly, the TiO_2_ Spin tips were conditioned with 20 µL of solution A (80% acetonitrile, 0,1% TFA), centrifuged at 3,000 × g for 2min and equilibrated with 20µL of solution B (75% acetonitrile, 0,075% TFA, 25% lactic acid) followed by centrifugation at 3,000 × g for 2 min. Peptides were resuspended in 10 µL of 2% TFA, mixed with 100 µL of solution B and centrifuged at 1,000 × g for 10min. Sample was applied back to the TiO_2_ Spin tips two more times in order to increase the adsorption of the phosphopeptides to the TiO_2_. Spin tips were washed with, sequentially, 20 µL of solution B and two times with 20 µL of solution A. Phosphopeptides were eluted by the sequential addition of 50 µL of 5% NH_4_OH and 50 µL of 5% pyrrolidine. Centrifugation was carried out at 1,000 × g for 5 min.

Phosphopeptides were further purified using GC Spin tips (GL-Tip, Titansphere, GL Sciences Inc, Japan). Briefly, the GC Spin tips were conditioned with 20 µL of solution A, centrifuged at 3,000 × g for 2 min and equilibrated with 20 µL of solution C (0,1% TFA in HPLC-grade water) followed by centrifugation at 3,000 × g for 2 min. Eluted phosphopeptides from the TiO_2_ Spin tips were added to the GC Spin tips and centrifuged at 1,000 × g for 5 min. GC Spin tips were washed with 20 µL of solution C. Phosphopeptides were eluted with 70 µL of solution A (1,000 × g for 3 min) and vacuum dried.

#### nanoLC-MS/MS protein identification and quantification

Samples were resuspended in 12 µL of 0.1% TFA in HPLC-grade water. For each run, 5 µL was injected in a nanoRSLC-Q Exactive PLUS (RSLC Ultimate 3000, Thermo Scientific, MA, USA). Phosphopeptides were loaded onto a µ-precolumn (Acclaim PepMap 100 C18, cartridge, 300 µm i.d.×5 mm, 5 µm, Thermo Scientific, MA, USA) and were separated on a 50 cm reversed-phase liquid chromatographic column (0.075 mm ID, Acclaim PepMap 100, C18, 2 µm, Thermo Scientific, MA, USA). Chromatography solvents were (A) 0.1% formic acid in water, and (B) 80% acetonitrile, 0.08% formic acid. Phosphopeptides were eluted from the column with the following gradient 5% to 40% B (180 min), 40% to 80% (6 min). At 181 min, the gradient returned to 5% to re-equilibrate the column for 20 minutes before the next injection. Two blanks were run between each sample to prevent sample carryover. Phosphopeptides eluting from the column were analyzed by data dependent MS/MS, using top-8 acquisition method. Phosphopeptides were fragmented using higher-energy collisional dissociation (HCD). Briefly, the instrument settings were as follows: resolution was set to 70,000 for MS scans and 17,500 for the data dependent MS/MS scans in order to increase speed. The MS AGC target was set to 1 × 10^6^ counts with maximum injection time set to 250 ms, while MS/MS AGC target was set to 2 × 10^5^ with maximum injection time set to 250 ms. The MS scan range was from 400 to 1.800 m/z. MS and MS/MS scans were recorded in profile mode. Dynamic exclusion was set to 30 seconds.

#### Data Processing Following nanoLC-MS/MS acquisition

The MS files were processed with the MaxQuant software version 1.5.3.30 and searched with Andromeda search engine against the UniProtKB/Swiss-Prot *F. tularensis* subsp. *novicida* database (release 28-04-2014, 1719 entries). To search parent mass and fragment ions, we set an initial mass deviation of 4.5 ppm and 0.5 Da respectively. The minimum peptide length was set to 7 amino acids and strict specificity for trypsin cleavage was required, allowing up to two missed cleavage sites. Carbamidomethylation (Cys) was set as fixed modification, whereas oxidation (Met), N-term acetylation and phosphorylation (Ser, Thr, Tyr) were set as variable modifications. The match between runs option was enabled with a match time window of 0.7 min and an alignment time window of 20 min. The false discovery rates (FDRs) at the protein and peptide level were set to 1%. Scores were calculated in MaxQuant as described previously(Cox & Mann, 2008). The reverse and common contaminants hits were removed from MaxQuant output.

The phosphopeptides output table and the corresponding logarithmic intensities were used for phosphopeptide analysis. The phosphopeptide table was expanded to separate individual phosphosites, and we kept all sites identified at least once in the three independent replicates of the analysis of *F. novicida*.

All data analysis of mass spectrometry data was performed in Perseus 1.6.0.7. For functional class annotation, we also integrated COG and EggNog databases in Perseus 1.6.0.7, freely available at www.perseus-framework.org. Data are available via ProteomeXchange with identifier PXD009225.

### Cell cultures and cell infection experiments

J774A.1 (ATCC^®^ TIB-67™) cells were propagated in Dulbecco’s Modified Eagle’s Medium (DMEM, PAA), containing 10% fetal bovine serum (FBS, PAA) unless otherwise stated. The day before infection, approximately 2×10^5^ eukaryotic cells per well were seeded in 12-well cell tissue plates and bacterial strains were grown overnight in 5 mL of Schaedler K3 at 37°C. Infections were realized at a multiplicity of infection (MOI) of 100 and incubated for 1 h at 37°C in culture medium. After 3 washes with cellular culture medium, plates were incubated for 4, 10 and 24 h in fresh medium supplemented with gentamycin (10 µg mL^−1^). At each kinetic point, cells were washed 3 times with culture medium and lysed by addition of 1 mL of distilled water for 10 min at 4°C. The titre of viable bacteria was determined by spreading preparations on chocolate plates. Each experiment was conducted at least twice in triplicates.

### Time lapse microscopy

J774-1 cells were transduced with the IncuCyte^®^ NucLight Red Lentivirus, following the manufacturer’s recommendations, to obtain red nuclear labelling of living cells due to the expression of the red fluorescent protein mKate2 containing a nuclear localization signal.

One prior to infection, J774-1 cells were seeded in 12-well cell tissue plates. Cells were infected (MOI of 1,000) with wild-type and mutant GFP-expressing cell. Synchronization of bacterial entry was realized by a 5 minutes centrifugation at 1,000 rpm. Plates were then incubated for 1 h at 37°C in culture medium. After 3 washes with cellular culture medium, plates were incubated at 5% CO_2_ and 37°C for 24 hours in fresh medium supplemented with gentamycin (10 µg mL^−1^). Bacterial multiplication was monitored in the fully automated microscope Incucyte^®^ S3 (Essen BioScience). Images were taken every 20 minutes with the 20X objective. Analysis and time-lapse videos (from which images were extracted) were generated by using Incucyte^®^ S3 software.

### Confocal experiments

J774.1 macrophage cells were infected (MOI of 1,000) with wild-type *F. novicida* U112, the isogenic Δ*iglB* mutant, the Δ*iglB* complemented either with wild-type *iglB* (CpWT) Y139A, Y139F mutants or an isogenic Δ*FPI* strain, in standard DMEM (DMEM-glucose) for 30 min at 37°C. Cells were then washed three times with PBS and maintained in fresh DMEM supplemented with gentamycin (10 μg mL^−1^) until the end of the experiment. Three kinetic points (*i.e.* 1 h, 4 h and 10 h) were sampled. For each time point, cells were washed with 1X PBS, fixed 15 min with 4% paraformaldheyde, and incubated 10 min in 50 mM NH_4_Cl in 1X PBS to quench free aldehydes. Cells were then blocked and permeabilised with PBS containing 0.1% saponin and 5% goat serum for 10 min at room temperature. Cells were then incubated for 30 min with anti-*F. novicida* mouse monoclonal antibody (1/500e final dilution, Creative Diagnostics) and anti-LAMP1 rabbit polyclonal antibody (1/100e, ABCAM). After washing, cells were incubated for 30 min with Alexa488-conjugated goat anti mouse and Alexa546 conjugated donkey anti rabbit secondary antibodies (1/400e, AbCam). After washing, DAPI was added (1/1,000) for 1 min and glass coverslips were mounted in Mowiol (Cityfluor Ltd.). Cells were examined using an X63 oil-immersion objective on a Zeiss Apotome 2 microscope. Co-localisation tests were quantified by using Image J software; and mean numbers were calculated on more than 500 cells for each condition. Confocal microscopy analyses were performed at the Cell Imaging Facility (Faculté de Médecine Necker Enfants-Malades).

### BMDMs Infections

Infection of WT or *ASC*^**-/-**^ BMDMs was performed as described previously (Rigard et al., 2016). Briefly, BMDMs were differentiated in DMEM (Invitrogen) with 10% v/v FCS (Thermo Fisher Scientific), 15% MCSF (L929 cell supernatant), 10 mM HEPES (Invitrogen), and non-essential amino acids (Invitrogen). One day before infection, macrophages were seeded into 12- 48- or 96-well plates at a density of 2×10^5^, 1.5×10^5^ or 5×10^4^ cells per well, respectively and incubated at 37°C, 5% CO_2_. The overnight culture of bacteria was added to the macrophages at multiplicity of infection (MOI) of 100. The plates were centrifuged for 15 min at 500 g to ensure comparable adhesion of the bacteria to the cells and placed at 37°C for 1h. After 3 washes with PBS, fresh medium with 5 µg mL^−1^ gentamycin (Invitrogen) was added to kill extracellular bacteria and plates were incubated for the desired time.

### Phagosomal rupture assay

Quantification of vacuolar escape using the β-lactamase/CCF4 assay (Life technologies) was performed as previously described(Meunier, Wallet et al.). *ASC*^**-/-**^ BMDMs seeded onto non-treated plates were infected as described above for 2 h, washed and incubated in CCF4 for 1 h at room temperature in the presence of 2.5 mM probenicid (Sigma). Propidium iodide negative cells were considered for the quantification of cells containing cytosolic *F. novicida* using excitation at 405 nm and detection at 450 nm (cleaved CCF4) or 510 nm (intact CCF4).

### Mouse infection

We evaluated the impact of the mutations Y139A and Y139F on bacterial virulence in the mouse model of tularemia, using an *in vivo* competition assay (Gesbert et al., 2014). Wild-type *F. novicida*, the Δ*iglB* mutant and the Δ*iglB* mutant complemented either with wild-type *iglB* (CpWT) or with Y139A and Y139F mutants, were grown in Schaedler K3 to exponential growth phase and diluted to the appropriate concentrations. 6 to 8-week-old female BALB/c mice (Janvier, Le Genest St Isle, France) were intraperitoneally (i.p.) inoculated with 200 μL of bacterial suspension. The actual number of viable bacteria in the inoculum was determined by plating dilutions of the bacterial suspension on chocolate plates. For competitive infections, wild-type *F. novicida* and mutant bacteria were mixed in 1:1 ratio and a total of 100 bacteria were used for infection of each of five mice. After two days, mice were sacrificed. Homogenized spleen and liver tissue from the five mice in one experiment were mixed, diluted and spread on to chocolate agar plates. Kanamycin selection to distinguish wild-type and mutant bacteria was performed. The competitive index (CI) was determined as the ratio (mutant output/WT output) / (mutant input/WT input). Statistical analysis for CI experiments was as described in (Brotcke, Weiss et al., 2006). Macrophage experiments were analyzed by using the Student’s unpaired t test.

### Sequence analyses

To retrieve IglB homologous proteins, we used the Hidden Markov Model (HMM) profile “T6SSii_iglB.hmm” from reference (Abby et al., 2016). We scanned a dataset of 2,462 predicted proteomes of complete bacterial genomes retrieved from GenBank Refseq (last accessed September 2016) using *hmmsearch* program of HMMER v.3.1b2 (gathering threshold = 25).

Our dataset included 19 genomes of the *Francisella* genus. All IglB sequences encoded by *Francisella* genomes were aligned with MUSCLE v.3.8.31 (Edgar, 2004). Due to differences in the annotation of the first methionine codon, we used the sequence of FTN_1323 as a reference for the multiple sequence alignment.

Redundancy of the sequence set retrieved from *hmmsearch* was reduced using cd-hit-v4.6.7 (Fu, Niu et al., 2012) with a 90% identity threshold. The longest representative sequence of each cluster was then aligned with MUSCLE v.3.8.31 (Edgar, 2004). Outliers that were very divergent in sequence length were removed. PROMALS3D and ESPript3.0 (Robert & Gouet, 2014) servers were used to visualize multiple sequence and structure alignment using 3J9O.B PDB structure reference file (Clemens et al., 2015). To determine the conservation of the phosphotyrosine site identified in *Francisella* genus (Tyr139 or Y139) we focused on the 12 amino acids surrounding the *Francisella* Tyr139 residue in the 535 aligned sequences.

To determine the conservation of the phosphotyrosine site identified in *Francisella* genus (Tyr139) we focused on the 12 amino acids surrounding the *Francisella* Y139 residue in the 535 aligned sequences. Out of the 535 aligned sequences, we found 90 sequences with a tyrosine residue in the vicinity of *Francisella* Tyr139 residue. For illustrative purpose, we selected 12 representative sequences out of 90 as shown in **Fig. S3**.

## Data Availability

The mass spectrometry proteomics data have been deposited to the ProteomeXchange Consortium via the PRIDE (Perez-Riverol, Xu et al., 2016) partner repository with the dataset identifier PXD012507.

## Acknowledgements

We thank Dr A. Sjostedt for providing the *Francisella* strain U112 and its *ptpA* transposon insertion derivative (*FTN_1046*) and Alain Schmitt (Cochin Institute Electron Microscopy Facility) for excellent technical support.

## Authors contribution

J.Z. performed most of the in vitro experiments; F.T. performed murine in vivo experiments and some immunofluorescence experiments; I.C.G and C.C. performed the proteomic analyses and I.C.G analyzed and compiled the data; N. K. performed the 3D analysis; A.J. performed the sequence alignments and analyzed the data ; H.R and M.C. performed the time lapse microscopy analyses; G.P. performed some in vitro experiments; M.B. and T.H. analyzed the data ; A.C. designed the experiments and J.Z. and A.C. analyzed the data, and wrote the paper.

The funders had no role in study design, data collection and analysis, decision to publish, or preparation of the manuscript.

## Supplementary materials

**Figure supplement 1. Schematic representation of the FPI locus and overview of the T6SS structure of *Francisella***

The *Francisella* FPI locus representation (*anmK* to *pdpA* gene) with nomenclature of the canonical T6SS and the *F. novicida* T6SS. A color was assigned for each gene function and applied on the overview of the T6SS structure. Red/Green/Blue/Orange, structural protein; Grey, secreted protein; White, hypothetical protein

**Figure supplement 2. Schematic representation of the genomic regions encoding proteins whose expression vary upon KCl stimulation.**

(**A**) Genomic organization of the FNI; (**B**). the Fe-S cluster- and iron uptake-associated loci. Genes colored in blue correspond to proteins present in decreased amounts; genes colored in rose, to proteins present in increased amounts, genes in white, unchanged.

**Figure supplement 3. Alignments of orthologous IglB proteins**

(**A**) Alignments of orthologous IglBs proteins showing conservation of Y139 in *Francisellae*. The IglB sequences retrieved from 19 *Francisella* genomes were aligned using MUSCLE (Edgar, 2004) and rendered using ESPript3.0 (Robert & Gouet, 2014). Secondary structure elements information is extracted from 3J9O PDB file (Clemens et al., 2015) and is displayed at the top of the alignment. Sequences are identified by corresponding species name followed by strain name and locus tag. Species names are abbreviated as follow: Fn, *Francisella tularensis* subsp. *novicida*; Ftm, *Francisella tularensis* subsp. *mediasiatica*; Fth, *Francisella tularensis* subsp. *holarctica*; Ftt, *Francisella tularensis* subsp. *tularensis*; Ft, *Francisella tularensis*; Fno, *Francisella noatunensis*; Fsp, *Francisella* sp. (**B**) Alignments of orthologous IglBs in other genera showing conservation of a tyrosine residue. Sequences of 11 representative IglB orthologs with a tyrosine residue in the vicinity of Y139 were aligned using MUSCLE (Edgar, 2004) and rendered using ESPript3.0 (Robert & Gouet, 2014). Secondary structure elements information is extracted from 3J9O PDB file (Clemens et al., 2015) and is displayed at the top of the alignment. Sequences are identified by corresponding Uniprot accession number. Species names are abbreviated as follow: FRATN, *Francisella tularensis subsp. novicida*; SALTY, *Salmonella* Typhimurium; YERPE, *Yersinia pestis;* BURMA, Burkholderia mallei; XANC5, Xanthomonas campestris; RALP1, Ralstonia pickettii; PHOAA, Photorhabdus asymbiotica; ERWAC, Erwinia amylovora; ENTCC, Enterobacter cloacae; BRAJP, Bradyrhizobium japonicum; RHIFR, Rhizobium fredii; 9GAMM, Serratia nematodiphila.

**Figure supplement 4. Effect of IglB mutations on growth in broth**

Wild-type *F. novicida* (WT, Grey squares), isogenic ∆*iglB* mutant (Δ*iglB*, blue triangles), and complemented *iglB* strain (Cp-WT, grey circles; Y139A, purple square; Y139F, red square), were grown in Schaedler K3 (left panel) or in Tryptic Soy broth (TSB) supplemented with 0,2% cysteine (right panel).

**Figure supplement 5. Intracellular multiplication in bone marrow-derived macrophages**

(**A**) Intracellular bacterial multiplication of wild-type *F. novicida* (WT, Grey squares), isogenic ∆*iglB* mutant (Δ*iglB*, blue triangles) and complemented Δ*iglB* strain in bone marrow-derived macrophages (BMDM) from BALB/c mice. (**B**) CCF4 assay. Infection of *ASC*^−/-^ BMDM were performed at MOI of 100 and the cytosol of BMDMs was loaded with CCF4, as described previously (Ramond et al., 2015). *F. novicida* naturally expresses a β-lactamase able to cleave the CCF4. Bacterial escape into the host cytosol is thus associated with a shift of the CCF4 probe fluorescence emission from 535 nm to 450 nm. The number of cells demonstrating CCF4 cleavage, determined at different time points (1 h, 3 h and 6 h) by flow cytometry, reached above 50% at 6 h for both WT and CpWT complemented strains whereas it was below 1% for the other mutant strains tested. (**C**) Groups of five female BALB/c mice were infected intraperitoneally with a mixture of 100 CFU of wild-type *F. novicida* and 100 CFU of Δ*iglB* mutant strain or complemented strain (Cp-WT, Y139A, Y139F). Bacterial burden was quantified in liver (L) and spleen (S) of mice. The data represent the competitive index (CI) value (in ordinate) for CFU of mutant/wild-type of each mouse, after 48 h infection, divided by CFU of mutant/wild-type in the inoculum. The CI recorded for the two *iglB* mutants was very low in both spleen and livers 2.5 days after infection (close to 10^−7^). As a control, we also performed an *in vivo* competition assay between wild-type and the *∆iglB* complemented strain (WT and CpWT). The competition index recorded close to 1 for CpWT (≤10^−1^ in both target organs), confirmed almost complete functional complementation.

Bars represent the geometric mean CI value.

**Figure supplement 6. Substitution of Tyr139 by phosphomimetics (Glu or Asp)**

(**A**) J774.1 macrophage-like cells were infected in DMEM with 100 MOI of wild-type *F. novicida* (WT) and mutated strains (Y139E, Y139D, Δ*FPI*) for 24 h. Results are shown as the average of log_10_ cfu mL^−1^ ± standard deviation. Each experiment was performed in triplicate. **, *p*<0.001 (as determined by two-tailed unpaired Student's *t*-test). (**B**) WB with anti-IglB of sucrose gradient fractions.

**Figure supplement 7. Inactivation of genes encoding proteins bearing a putative tyrosine kinase domain**

J774.1 macrophage-like cells were infected in DMEM-High glucose with an MOI of 100 with wild-type *F. novicida* (WT), three mutants corresponding to a transposon insertion into genes *FTN_1082*, *FTN_1433*, *FTN_0459* (tnfn1_pw060323p01q162, tnfn1_pw060420p02q142, tnfn1_pw060323p07q132, respectively from the BEI resources repository), here designated Δ*1082*, Δ*1433*, Δ*0459,* for simplification), and a Δ*FPI* mutant (Δ*FPI*). Intracellular multiplication was monitored at 10 h. Results are shown as the average of log10 cfu mL^−1^ ± standard deviation. Each experiment was performed in triplicate.

**Video supplement 1.** WT: J774.1 macrophage-like cells were infected in DMEM-High glucose at an MOI of 1,000 with wild-type *F. novicida* ∆*iglB* expressing pKK214::pGro*iglB*-pGro*gfp* (designated cpWT-GFP).

**Video supplement 2.** Y139F: J774.1 macrophage-like cells were infected in DMEM-High glucose at an MOI of 1,000 with wild-type *F. novicida* ∆*iglB* expressing pKK214::pGro*iglB*Y139F-pGro*gfp* (designated cpY139F-GFP)

**Table supplement 1. Whole cell proteome of non-induced and KCl-induced *F. novicida.*** In this table, we report the log2 (intensity) of the proteins identified in whole cell fractions from non-induced and KCl-induced wild-type *F. novicida*. Peptides and sequence coverage indicates the number of total peptides identified and the primary sequence coverage for that protein across samples. The Q value and the score refer to the quality of the identification.

**Table supplement 2. Detailed list of phosphosites identified in *F. novicida***

In this table, we report: i) the average of the log2 (intensity) of the phosphosites identified from three independent replicates; ii) the amino acid carrying the phosphorylation (“Amino Acid”); iii) its position on the protein (Position); iv) the “localization probability” of the phosphosite (1 being 100%); v) the number of replicates the phosphosite has been identified in (“identified”); vi) the “sequence window” of 15 amino acids before and after the amino acid phosphorylated; vii) the probability and score of identification and localization (PEP, score, score for localization); viii) the “charge” of the peptide; and ix) the annotation according to COG and EggNOG.

**Table supplement 3. Table of intensities for IglB peptide carrying Y139 phosphorylation.**

**Table supplement 4. List of proteins identified in the “Heavy” (H) and “Light” (L) Fraction in WT and in the Y139F mutant**

WT_H, WT_L, Y139_H, Y139F_L report the log2 (intensity) of the proteins identified in the fractions. Peptides and sequence coverage indicates the number of total peptides identified and the primary sequence coverage for that protein across samples. The Q value and the score refer to the quality of the identification.

**Table supplement 5. Strains and plasmids.**

**Table supplement 6. Primers.**

